# Compositional analysis of ALS-linked stress granule-like structures reveals factors and cellular pathways dysregulated by mutant FUS under stress

**DOI:** 10.1101/2021.03.02.433611

**Authors:** Haiyan An, Gioana Litscher, Wenbin Wei, Naruaki Watanabe, Tadafumi Hashimoto, Takeshi Iwatsubo, Vladimir L. Buchman, Tatyana A. Shelkovnikova

**Author notes:** To whom correspondence should be addressed at Medicines Discovery Institute, Cardiff University, Park Place, Cardiff, CF10 3AT, United Kingdom.

## Abstract

Formation of cytoplasmic RNA-protein structures called stress granules (SGs) is a highly conserved cellular response to stress. Abnormal metabolism of SGs may contribute to the pathogenesis of (neuro)degenerative diseases such as amyotrophic lateral sclerosis (ALS). Many SG proteins are affected by mutations causative of these conditions, including fused in sarcoma (FUS). Mutant FUS variants have high affinity to SGs and also spontaneously form *de novo* cytoplasmic RNA granules. Mutant FUS-containing assemblies (mFAs), often called “pathological SGs”, are proposed to play a role in ALS-FUS pathogenesis. However, global structural differences between mFAs and physiological SGs remain largely unknown, therefore it is unclear whether and how mFAs may affect cellular stress responses. Here we used affinity purification to characterise the protein and RNA composition of normal SGs and mFAs purified from stressed cells. Comparison of the SG and mFA proteomes revealed that proteasome subunits and certain nucleocytoplasmic transport factors are depleted from mFAs, whereas translation elongation, mRNA surveillance and splicing factors as well as mitochondrial proteins are enriched in mFAs, as compared to SGs. Validation experiments for a hit from our analysis, a splicing factor hnRNPA3, confirmed its RNA-dependent sequestration into mFAs in cells and into pathological FUS inclusions in a FUS transgenic mouse model. Furthermore, silencing of the *Drosophila* hnRNPA3 ortholog dramatically enhanced FUS toxicity in transgenic flies. Comparative transcriptomic analysis of SGs and mFAs revealed that mFAs recruit a significantly less diverse spectrum of RNAs, including reduced recruitment of transcripts encoding proteins involved in protein translation, DNA damage response, and apoptotic signalling. However mFAs abnormally sequester certain mRNAs encoding proteins involved in stress signalling cascades. Overall, our study establishes molecular differences between physiological SGs and mFAs and identifies the spectrum of proteins, RNAs and respective cellular pathways affected by mFAs in stressed cells. In conclusion, we show that mFAs are compositionally distinct from SGs and that they cannot fully substitute for SG functions while gaining novel, potentially toxic functions in cellular stress response. Results of our study support a pathogenic role for stress-induced cytoplasmic FUS assemblies in ALS-FUS.

## Introduction

Amyotrophic lateral sclerosis (ALS) is a rapidly progressive, incurable and inevitably fatal neuromuscular disease affecting motor neurons in the spinal cord and motor cortex. Up to 90% of ALS cases are sporadic (sALS), and 10% are caused by mutations in known genes (familial ALS, fALS) (Hardiman et al., 2017). Despite recent genetic and molecular breakthroughs in dissecting ALS pathogenesis, underlying mechanisms shared by fALS and sALS are poorly understood, which represents a major obstacle in establishing therapeutic targets for this devastating disease (Taylor et al., 2016).

Stress granules (SGs) are cytoplasmic RNA granules that form as a normal cellular response to stresses involving a shutdown of protein translation. SGs may serve to shield RNA from degradation until protein translation can be safely resumed (Kedersha and Anderson, 2002) and to modulate stress signalling, including anti-apoptotic signalling, selective translation of molecular chaperones and adjustment of protein translation rates to prevent the accumulation of misfolded proteins (Kedersha et al., 2013).

Multiple SG proteins are affected by ALS-causative mutations (Li et al., 2013), including fused in sarcoma, or FUS (Kwiatkowski et al., 2009; Vance et al., 2009). In the majority of ALS-FUS cases, the nuclear localisation signal (NLS) of FUS protein bears single amino acid substitutions or deletions, causing a defect in its nuclear import, uncontrollable deposition and inclusion formation in the cytoplasm – FUS proteinopathy (Lattante et al., 2013; Mackenzie et al., 2010). Unlike normal protein, mutant FUS isoforms mislocalised to the cytoplasm are readily recruited into stress-induced SGs (Bosco et al., 2010; Dormann et al., 2010). In addition, as we and others showed, overexpressed or endogenous mutant FUS spontaneously forms cytoplasmic microaggregates that represent a novel type of RNP granule we termed the FUS granule (Japtok et al., 2015; Kino et al., 2015; Shelkovnikova et al., 2019; Shelkovnikova et al., 2014a; Takanashi and Yamaguchi, 2014). In stressed cells, FUS granules can coalesce into larger assemblies comparable in size with mature SGs – FUS aggregates. Presence of FUS aggregates is generally disruptive for physiological SGs because these aberrant structures compete with SGs for their core proteins such as G3BP1 and TIAR, as well as RNA species serving as a scaffold for SGs (Shelkovnikova et al., 2014a). Similarly, enrichment of mutant FUS in SGs can alter their dynamics (Aulas and Vande Velde, 2015), interactions between components and hence SG function. However, the relationships between normal SGs and pathological FUS-containing SG-like structures remain unclear. For example, it is possible that the latter structures can at least partially substitute for normal SG function during stress however they are unlikely to adopt the full range of SG roles. FUS-containing SG-like structures can also gain novel unwanted (toxic) functions. Stress likely plays a pivotal role as a secondary trigger of ALS-causative pathology (Al-Chalabi et al., 2014; Dormann et al., 2010; Shelkovnikova et al., 2019), therefore understanding structural and functional differences between normal SGs and pathological, mutant FUS-containing assemblies in stressed cells can provide us with important clues on mutant FUS-induced dysregulation of stress signalling and its contribution to disease processes in ALS. Furthermore, while strong experimental evidence supports the role of cytoplasmic gain of FUS function in ALS-FUS pathogenesis (Devoy et al., 2017; Scekic-Zahirovic et al., 2016; Sharma et al., 2016), the contribution of cytoplasmic FUS aggregation into the pathology development is still debated.

In the current study, we interrogated the protein and RNA composition of biochemically purified mutant FUS cytoplasmic assemblies (mutant FUS containing SGs and FAs combined) – “mFAs”, and compared their proteome and transcriptome to those of physiological SGs purified in parallel. This approach enabled identification of a range of protein and RNA species enriched or depleted in mFAs, as compared to physiological SGs, and hence respective pathways impacted by mFA presence in the cytoplasm of stressed cells. In particular, our analysis highlighted possible deficiencies in proteasomal degradation of mFAs. In addition to that, we found that mFAs display enrichment in splicing, mitochondrial, protein translation and RNA quality control factors, as compared to SGs. Focusing on a hit from our proteomic analysis, hnRNPA3, an RNA-binding protein recently implicated in ALS with *C9ORF72* repeat expansions (Mori et al., 2013), we demonstrated that it interacts with FUS in an RNA-dependent manner and is sequestered into mFAs in cells and into FUS inclusions in transgenic mice. Furthermore, we found that depletion of its ortholog in a *Drosophila* model of ALS-FUS exacerbates FUS toxicity. At the transcript level, we revealed that mFAs fail to sequester some mRNAs encoding proteins involved in protein translation, DNA damage response, microtubule organisation and stress/apoptotic signalling, which are recruited into physiological SGs. On the other hand, some transcripts encoding stress-relevant proteins were found to be abnormally retained in mFAs.

Therefore, formation of cytoplasmic FUS assemblies is expected to have a profound effect on the protein and RNA homeostasis in neurons. Results of our study establish molecular differences between pathological, ALS-linked RNP granules and physiological SGs, and provide evidence in support of the role of cytoplasmic FUS aggregates in ALS-FUS pathogenesis.

## Results

### Purification of stress granule and mutant FUS assembly cores for comparative analysis of their protein and RNA composition

SGs were reported to consist of a denser “core” and a more diffuse “shell”, and relatively stable SG cores can be isolated from mammalian cells (Jain et al., 2016). Mammalian SG cores measured based on G3BP1-GFP fluorescence are ∼200 nm (233.1±18.6 nm) in diameter (Jain et al., 2016). This is close to the estimated size of spontaneous FUS granules formed by overexpressed or endogenous FUS – ∼150-200 nm (Shelkovnikova et al., 2019; Shelkovnikova et al., 2014a). Therefore, we reasoned that FUS granules can be enriched and purified using the protocol developed for SG cores. However, only ∼50% of mutant FUS-expressing cells develop FUS granules or FUS aggregates under basal conditions, whilst the remaining cells contain diffuse cytoplasmic FUS which becomes incorporated into SGs during stress (Shelkovnikova et al., 2014a). Therefore, in mutant FUS-expressing cells subjected to a SG-inducing stress, the SG core purification protocol will result in isolation of both FUS granules (as part of FUS aggregates) and mutant FUS-containing SG cores. This heterogeneous aggregated FUS fraction is thereafter referred as “mutant FUS assembly cores”, or “mFA cores”. We selected FUS with an ALS-associated point mutation R522G with confirmed ability to mislocalize to the cytoplasm and form FUS granules (Shelkovnikova et al., 2014a).

HEK293 cells were transfected to express either G3BP1-GFP or FUS(R522G)-GFP, and after 24 h, cells were treated with an oxidative stressor NaAsO_2_ (sodium arsenite) for 1 h. SG and mFA formation was confirmed in these cultures by fluorescent microscopy (Fig. 1A). Enriched SG and mFA cores were immunoprecipitated from cell lysates using GFP-Trap® beads, and final bead fractions were used for proteomic and transcriptomic analyses. Experimental pipeline is schematically shown in Fig. 1B. Successful SG and mFA core isolation was confirmed by western blot with an anti-GFP antibody (Fig. 1C).

**Figure 1.**
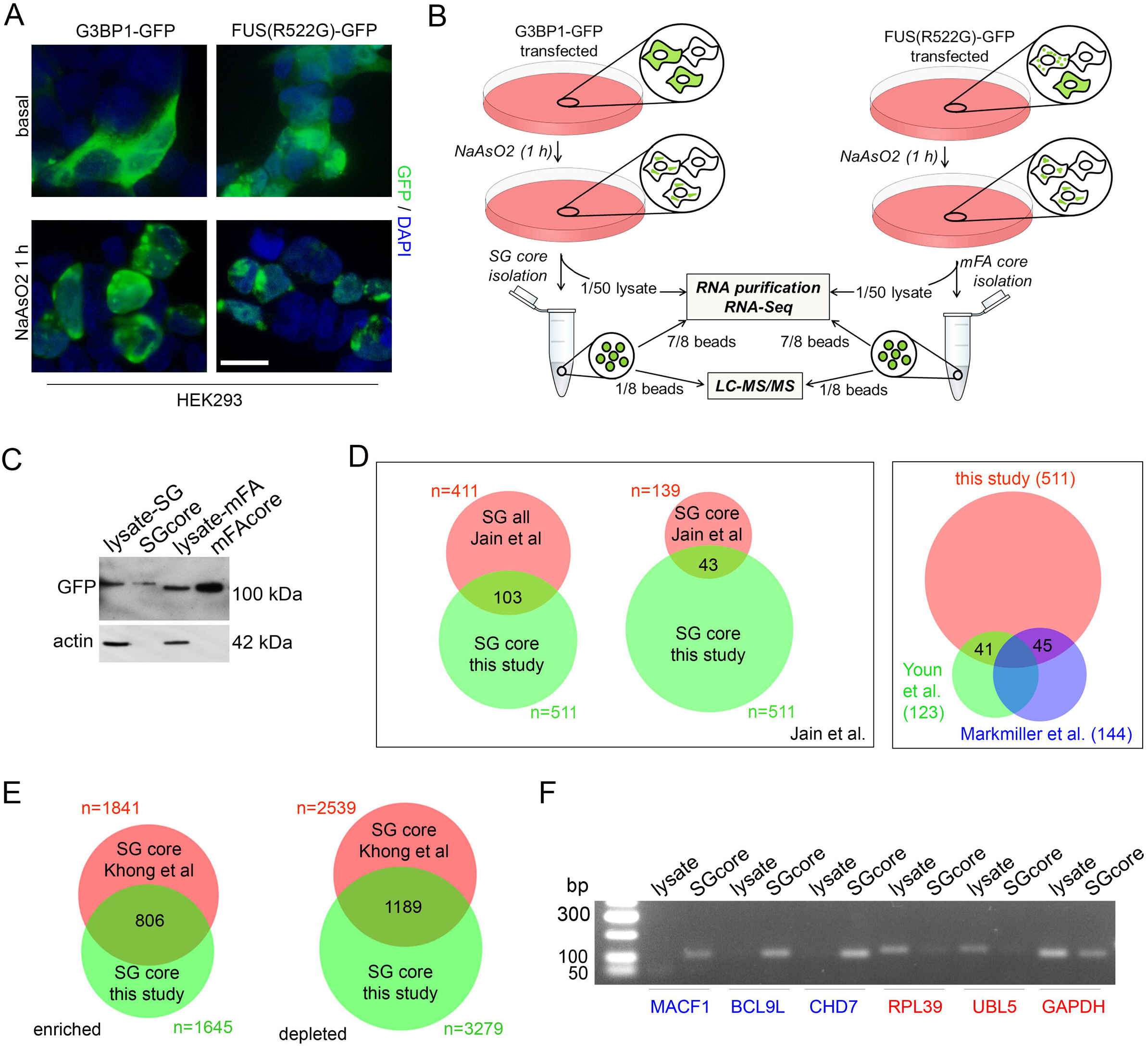
Parallel purification and quality control of SG and mFA cores. (A) Distribution of G3BP1-GFP and FUS(R522G)-GFP in HEK293 cells under basal conditions and under stress. Cells were transfected to express GFP-tagged proteins and 24 h post-transfection, treated with 0.5 mM NaAsO_2_ for 1 h. Scale bar, 10 µm. (B) Experimental pipeline for isolation of SG and mFA cores for proteomic and transcriptomic analyses. (C) Efficiency of SG and mFA core pull-down analyzed by western blot with an anti-GFP antibody. 2% of the respective lysate was loaded in each case. (D) Overlaps between the HEK293 SG_core_ proteomic dataset from the current study and published SG datasets. (E) Overlaps between the HEK293 (this study) and U2OS (Khong et al., 2017) SG_core_ transcriptomic datasets. (F) Recruitment of transcripts classified as enriched in SG cores (blue) or depleted from SG cores (red) in U2OS cells (Khong et al., 2017) into HEK293 SG cores, as analyzed by non-saturated RT-PCR.

Purified SG cores, mFA cores and the beads fraction from control (GFP only) samples were subjected to LC-MS/MS analysis. The “background” list of proteins obtained in “GFP only” samples was subtracted from SG_core_ and mFA_core_ protein lists, yielding final SG_core_ and mFA_core_ proteomes (Table S1). To ascertain successful purification of SG cores, we run comparisons of our SG_core_ proteome with the published SG proteomes. Our dataset demonstrated significant overlaps with the datasets from the original study (Jain et al., 2016); it was found to contain one-third (43/139) of the U2OS SG_core_ proteome (p-value of overlap 1.595e^-32^) and a quarter (103/411) of proteins from the “full SG proteome” compiled in the same study (p-value of overlap 8.950e^-68^) (Fig. 1D). One explanation for a larger size of our SG_core_ dataset and the partial overlap with the U2OS SG_core_ dataset is cell-specific differences. Consistently, one-third (41/123) of SG proteins from HEK293 cells identified by an APEX labeling method (Markmiller et al., 2018) were also included in our SG_core_ dataset (p-value of overlap 1.540e^-32^) (Fig. 1D). Our SG_core_ dataset also showed substantial enrichment in SG and P-body proteins from the HEK293 dataset obtained using proximity-dependent biotinylation (Youn et al., 2018) (45/144 proteins, p-value of overlap 3.261e^-34^). It should be noted that individual overlaps between our SG_core_ dataset and published datasets (43, 41 and 45 proteins for Jain et al., Markmiller et al. and Youn et al., respectively) do not correspond to the same list of proteins, as only 15 proteins were found to be common among the three datasets (Fig. 1D). Enrichr analysis on the overrepresented Biological Process GO terms showed a significant overlap between HEK293 and U2OS SG_core_ datasets (Fig. S1A). The top significant Biological Process GO terms in both datasets were related to regulation of translation (Fig. S1B).

RNA-Seq analysis was performed on the original cell lysates and separately on the final bead fractions containing SG/mFA cores (Fig. 1A). Differential gene expression analysis was used to determine the enrichment or depletion of RNAs in SG and mFA cores, as compared to the respective cell lysates (Table S2). We found that in total, 1,645 RNAs were enriched and 3,279 were depleted from SG cores in HEK293 cells (cut-off of p<0.05 and fold change of 1.2, Table S2). We next compared our HEK293 SG_core_ transcriptomic datasets with those from U2OS SG cores (Khong et al., 2017). Approximately one-third of HEK293 SG_core_-enriched and -depleted transcriptomes overlapped with those from U2OS cells (806/1,645 and 1,189/3,279, respectively) (p-value of overlap 4.792e^-322^) (Fig. 1E). Six out of ten most enriched cytosolic mRNAs in U2OS SG cores were included in the top 100 HEK293 SG_core_-enriched mRNAs (MACF1, AHNAK2, MAP1A, SPEN, BCL9L, KMT2D). Six transcripts (three enriched and three depleted) were validated by semi-quantitative PCR (Fig. 1F). Enrichr analysis on the overrepresented GO terms in HEK293 SG cores showed that they significantly overlap with those for the U2OS SG_core_ dataset (Fig. S1C). In particular, the top Biological Process GO terms from HEK293 and U2OS datasets (41 and 58 terms, respectively, adjusted p-value between 0.004 and 0.0009) were related to regulation of transcription and protein phosphorylation (Fig. S1D).

Thus, comparisons of our HEK293 SG_core_ proteome and transcriptome with the published data confirmed that our modified affinity purification protocol allows efficient isolation of SG core-like RNP granules.

### Pathways dysregulated by selective enrichment or depletion of specific proteins in mutant FUS assemblies during stress

Comparison of the SG_core_ and mFA_core_ protein datasets (n=511 and 457, respectively) obtained in parallel revealed a substantial overlap between the two: 238 proteins, or nearly half for both datasets, were found to be in common (Fig. 2A). We used the STRING v11 database in order to reconstruct and compare the networks of protein-protein interactions for the two types of RNP granules. Overall, mFA cores formed a tighter, more compact network of interactors as compared to SG cores (Fig. 2B). Among the top 25 enriched Biological Process GO term categories in the STRING database, five were shared by SG cores and mFA cores, namely *posttranscriptional regulation of gene expression, regulation of translation, mRNA metabolic process, amide biosynthetic process* and *peptide metabolic process*. We next analyzed common set of 238 genes using Enrichr. The top enriched Biological process GO term category for this list of proteins was *regulation of translation*, whereas the top Cellular Component GO term categories included *cytoplasmic ribonucleoprotein granules, cytoplasmic stress granules* and *P-bodies* (Fig. 2C).

**Figure 2.**
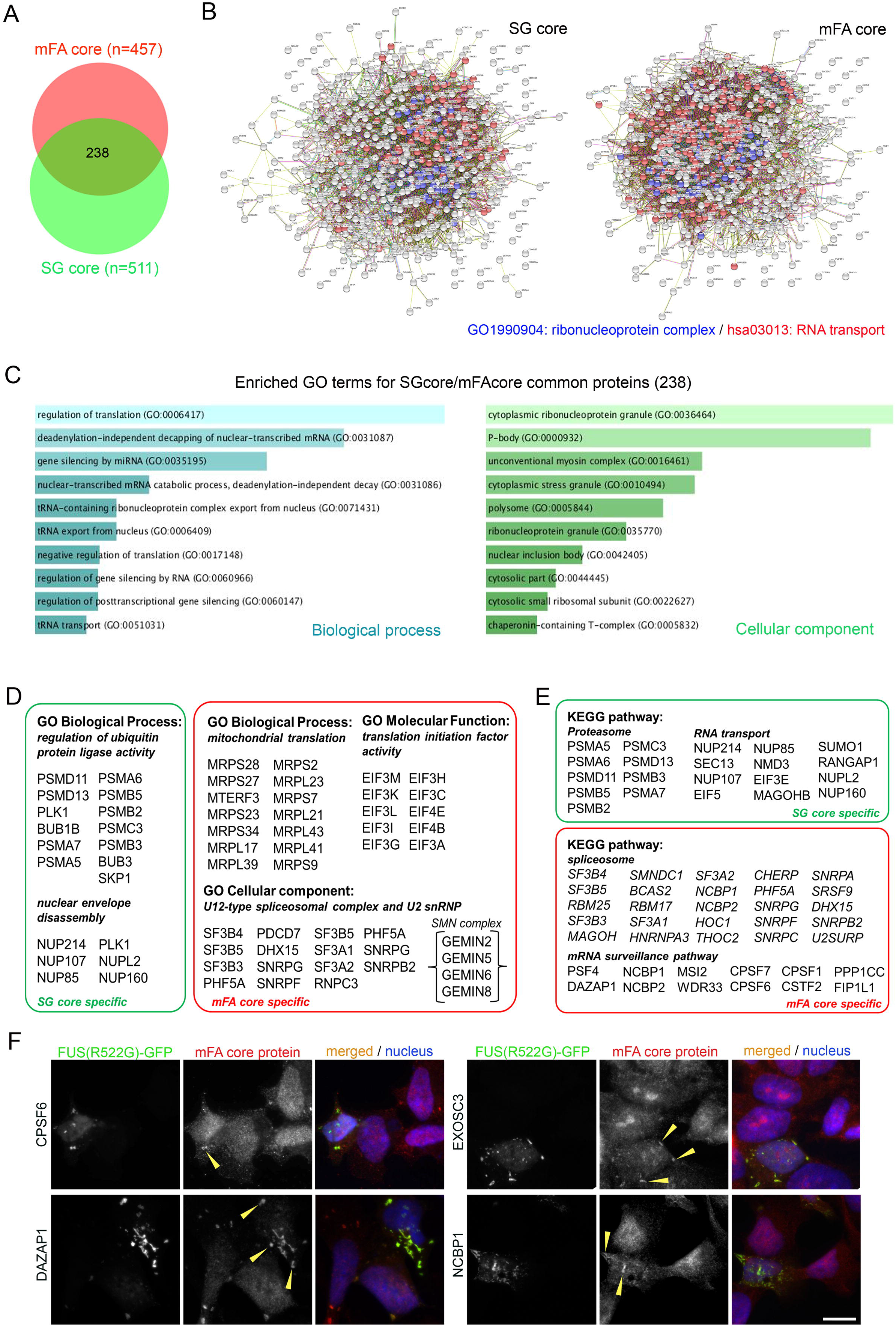
Comparative analysis of the proteomes of SG and FA cores. (A) Overlaps between the HEK293 SG_core_ and mFA_core_ proteomic datasets. (B) Protein networks for SG and mFA cores visualized using the STRING v.11 database. Proteins from the enriched categories: *Cellular component GO1990904: ribonucleoprotein complex* (FDR of 1.29^-28^ and 4.51^-85^ for SG and mFA cores, respectively) and *KEGG pathway hsa03013: RNA transport* (FDR of 1.86^-32^ and 1.66^-26^, for SG and mFA cores, respectively) are given in red and blue, respectively. (C) Overrepresented Biological Process and Cellular component GO terms for proteins shared by SG and mFA cores, as determined using the Enrichr tool. (D) GO terms overrepresented in the SG_core_-specific (n=273) and mFA_core_-specific (n=219) proteomic datasets. (E) KEGG pathways overrepresented in the SG_core_-specific (n=273) and mFA_core_-specific (n=219) proteomic datasets. (F) Validation of mFA_core_-recruited proteins. Presence of four proteins identified as components of mFAs formed by FUS(R522G)-GFP was verified by immunocytochemistry in SH-SY5Y cells. Cells were treated with 0.5 mM NaAsO_2_ for 1 h. Representative images are shown. Scale bar, 10 µm.

We next focused on the SG_core_- and mFA_core_-specific sets of proteins. Using Enrichr and subsequently Revigo to remove redundant GO terms, we analyzed the list of 273 proteins found in SG cores but not mFA cores. Non-redundant Biological Process GO terms returned for this dataset included *regulation of ubiquitin protein ligase activity; protein sumoylation; cellular response to hypoxia; nuclear envelope disassembly;* and *regulation of gene silencing by RNA* (cut-off of false discovery rate, FDR < 0.01). In particular, we found that multiple proteasome subunit proteins (PSMA7, PSMA5, PSMA6, PSMB5, PSMB2, PSMC3, PSMB3) were recruited into SG cores, whereas only one proteasome subunit protein, PSMA4, was present in the mFA_core_ dataset (Fig. 2D). The second category of proteins uniquely recruited into SG cores were nuclear envelope proteins including NUP214, NUP107, NUP85, NUPL2 and NUP160 (Fig. 2D). In total, nine nuclear pore complex (NPC) proteins were found to be enriched in SG cores, and only five – in mFA cores. Consistently, KEGG pathway analysis also highlighted *Proteasome* and *RNA transport pathways* as enriched in proteins identified specifically in SG cores (Fig. 2E).

Capture and detainment of proteins by mFAs may result in their loss of function. Analysis of the mFA_core_-specific proteome (n=219) showed significant enrichment of Biological Process GO terms related to mitochondrial metabolism and more specifically, mitochondrial translation (Fig. 2D). Secondly, we found significant enrichment of translation initiation factors in mFA cores, including multiple members of the eIF-3 complex (EIF3M, EIF3K, EIF3L, EIF3I, EIF3G, EIF3H, EIF3C, EIF3A). Another significantly enriched group of proteins in mFA cores were proteins involved in major (U2) and minor (U12) spliceosome subunits. In addition, we found that five Gemin proteins which interact with snRNPs to form the SMN complex are present in the mFA_core_ dataset, as opposed to only one in the SG_core_ dataset (Fig. 2D).

Finally, proteins within the *Spliceosome* and *mRNA surveillance pathway* KEGG pathways were significantly enriched in the FA_core_ dataset (Fig. 2E). Both included a number of multifunctional proteins with critical role in RNA metabolism, such as NCBP1, NCBP2, HNRNPA3, SRSF9 (*Spliceosome*); and DAZAP1, CPSF7, CPSF6, CPSF1, EXOSC3 (*mRNA surveillance pathway*). We confirmed the recruitment of CPSF6, DAZAP1, NCPB1 and EXOSC3 into mFAs in stressed cells by immunocytochemistry (Fig. 2F). It should be noted that CPSF6 and DAZAP1 were previously detected in physiological SGs by immunocytochemistry (An et al., 2019b), but not in SG cores by proteomics (Jain et al., 2016), therefore they are likely SG shell components.

### RNA transcripts differentially recruited into stress granules and mutant FUS assemblies

Comparative analysis of the SG_core_ and mFA_core_ transcriptomes revealed that significantly fewer RNAs were accumulated in mFA cores as compared to SG cores (802 *vs*. 1,645) suggesting that mFA cores are able to sequester a less diverse spectrum of transcripts. Overall, approximately two-thirds of SG_core_ transcripts (1,243 out of 1,645) failed to be efficiently recruited into mFA cores (Fig. 3A). In contrast, the overlap between RNA datasets depleted from SG cores and mFA cores was more prominent: 71.4% of mFA_core_-depleted transcripts were also included in the SG_core_ dataset (Fig. 3A). For example, two transcripts known to be enriched in SG cores, MACF and CHD7 (see Fig 1F), did not appear among mFA_core_-enriched transcripts, and BCL9L was found to be enriched in mFA cores 2.08-fold, as compared to 4.14-fold enrichment in the SG core (Table S2). Consistent with a larger overlap between SG_core_- and mFA_core_-excluded transcripts, RNAs depleted from SG cores (RPL39, UBL5, GAPDH) were also found to be depleted from mFA cores however depletion from SG cores was slightly more pronounced (Table S2, Fig. 3B). Furthermore, we found that on average, transcripts were significantly less enriched in mFA cores and that depletion of transcripts was also less dramatic in mFA cores, as compared to SG cores (Fig. 3C).

**Figure 3.**
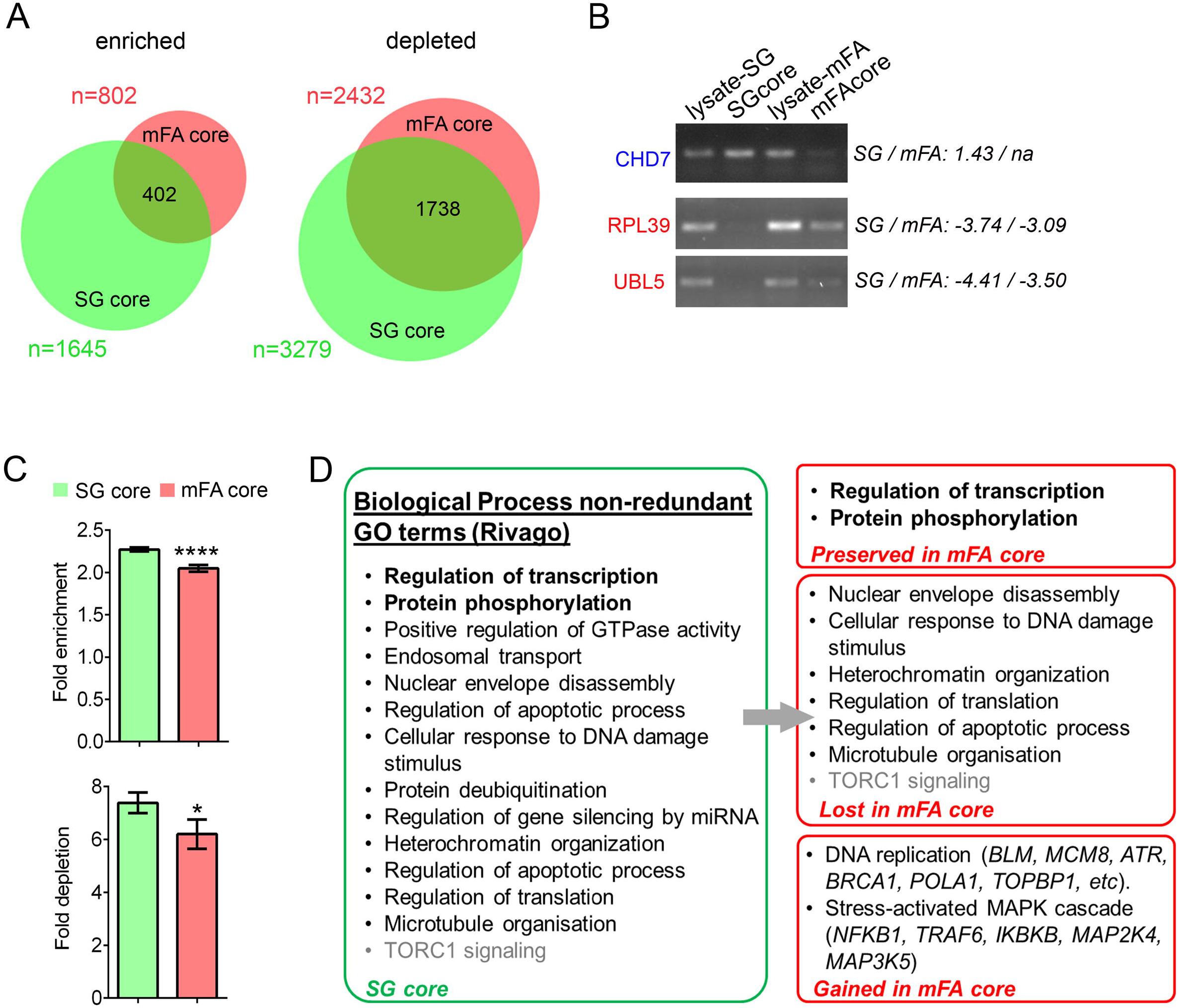
Comparative analysis of the transcriptomes of SG and mFA cores. (A) Overlaps between the HEK293 SG_core_ and mFA_core_ transcriptomic datasets. (B) Validation of transcripts enriched and depleted in mFA cores by non-saturated RT-PCR. Fold enrichment/depletion is also shown (na – not available, transcript did not show enrichment). (C) Attenuated enrichment and depletion of RNAs in mFA cores as compared to SG cores. *p<0.05, ****p<0.0001. Mean fold enrichment or depletion for the entire “enriched” and “depleted” SG and mFA datasets were used for analysis (see Table S2). (D) List of non-redundant Biological process GO terms overrepresented for the HEK293 SG_core_ dataset (identified in the full SG_core_ dataset using Enrichr and filtered using Rivago) (left) and list of GO terms preserved or lost in the mFA_core_ dataset. GO terms overrepresented in mFA cores dataset only (“Gained in mFA core”), together with examples of proteins from this GO term, are also shown.

We next performed GO term enrichment analysis on the mFA_core_ dataset, to be able to identify functional classes of transcripts over- and underrepresented in mFAs as opposed to physiological SGs. Two Biological process GO term categories, “Protein phosphorylation (GO:0006468)” and “Regulation of transcription, DNA-templated (GO:0006355)”, were found to be in common between SG and mFA cores (Fig. 3D). However, the number of RNAs in both categories was significantly lower in mFA cores: 53 *vs*. 89 out of 470 transcripts for *protein phosphorylation* and 105 *vs*. 243 out of 1,598 transcripts for *regulation of transcription* were present in mFA cores *vs*. SG cores. Furthermore, enrichment for some GO terms was lost in the mFA_core_ dataset, including those related to *nuclear envelope disassembly, cellular response to DNA damage stimulus, heterochromatin organization, regulation of apoptotic process, microtubule organization* (Fig. 3D). This was also due to dramatically decreased number of RNAs for mFA cores in these GO term categories, e.g. 8 *vs*. 38 out of 813 for *regulation of translation* and 48 *vs*. 94 out of 815 for *regulation of apoptotic process* for mFA cores *vs*. SG cores. Two GO term categories were found to be significantly overrepresented in the mFA_core_ but not in SG_core_ dataset, “DNA replication (GO:0006260)” and “Stress-activated MAPK cascade (GO:0051403)”. In particular, RNAs encoding the major proteins involved in MAPK signaling, NFKB1, TRAF6, IKBKB, MAP2K4, and MAP3K5, were found to be recruited into mFA cores but not SG cores (Fig. 3D).

Finally, we analyzed SG_core_- and FA_core_-specific transcripts (n=1,243 and n=400, respectively) to identify additional groups of transcripts selectively recruited to SG cores or mFA cores. Transcripts encoding proteins critical for TORC1 signaling (GO:0038202: MTOR, RPTOR, LARP1); heterochromatin organization (GO:0070828, DNMT1, BRCA1, BCOR, JARID2 and SUPT6H); regulation of microtubule polymerization (GO:0031113, DCNT1, MAPT); SWI/SNF complex (GO:0016514, SMARCC1, SMARCC2, SMARCA4) as well as disease-relevant AGO2 and ATM proteins, were found to be recruited to SG cores but not mFA cores. On the other hand, RNAs encoding JAK1, JAK2, MBP and all three TGF-beta cytokines (TGFB1, TGFB2 and TGFB3) were identified as enriched in mFA cores but not in SG cores (Fig. 3D).

### hnRNPA3 is sequestered into mutant FUS aggregates in vitro and in vivo and its depletion exacerbates FUS toxicity in Drosophila

We next focused on one of the hits from our proteomic analysis relevant to ALS pathogenesis, hnRNPA3. hnRNPA3 has been identified as the C9ORF72 repeat RNA interactor and a component of C9ORF72 dipeptide repeat inclusions that regulates repeat RNA levels; its loss of function might contribute to ALS-C9 pathogenesis (Mori et al., 2013; Nihei et al., 2020). We performed immunostaining of SGs and mFAs in stressed SH-SY5Y cells with an anti-hnRNPA3 antibody. SH-SY5Y cells were chosen over HEK293 cells since they are flatter cells with larger cytoplasm more suitable for imaging; it is also a cell line possessing a number of neuronal characteristics. Immunostaining demonstrated significant enrichment of endogenous hnRNPA3 in NaAsO_2_-induced mFAs formed by FUS(R522G)-GFP while it was virtually absent in SGs formed in G3BP-GFP expressing or untransfected cells (Fig. 4A). Co-expression of Flag-tagged hnRNPA3 and FUS(R522G)-GFP confirmed co-deposition of the two proteins (Fig. 4B). To establish FUS protein domains responsible for its interaction with hnRNPA3 and its recruitment into mFAs, we transiently expressed FUS deletion mutants localized to the cytoplasm (Fig. 4C), followed by anti-hnRNPA3 staining. This analysis showed that RRM and RGG domains of FUS are required for efficient recruitment of hnRNPA3 into mFAs (Fig. 4D) and suggested that FUS-hnRNPA3 interaction is RNA-dependent. To address this directly, we performed immunoprecipitation (IP) of mutant FUS from samples treated or not treated with RNase A, and examined the presence of endogenous hnRNPA3 in IP samples by western blot. We included WT FUS in this experiment to test whether a mutation can increase FUS affinity to hnRNPA3. WT and mutant (R522G) FUS precipitated hnRNPA3 with equal efficiency, and RNase A treatment completely abolished FUS-hnRNPA3 interaction both with normal and mutant FUS (Fig. 4E).

**Figure 4.**
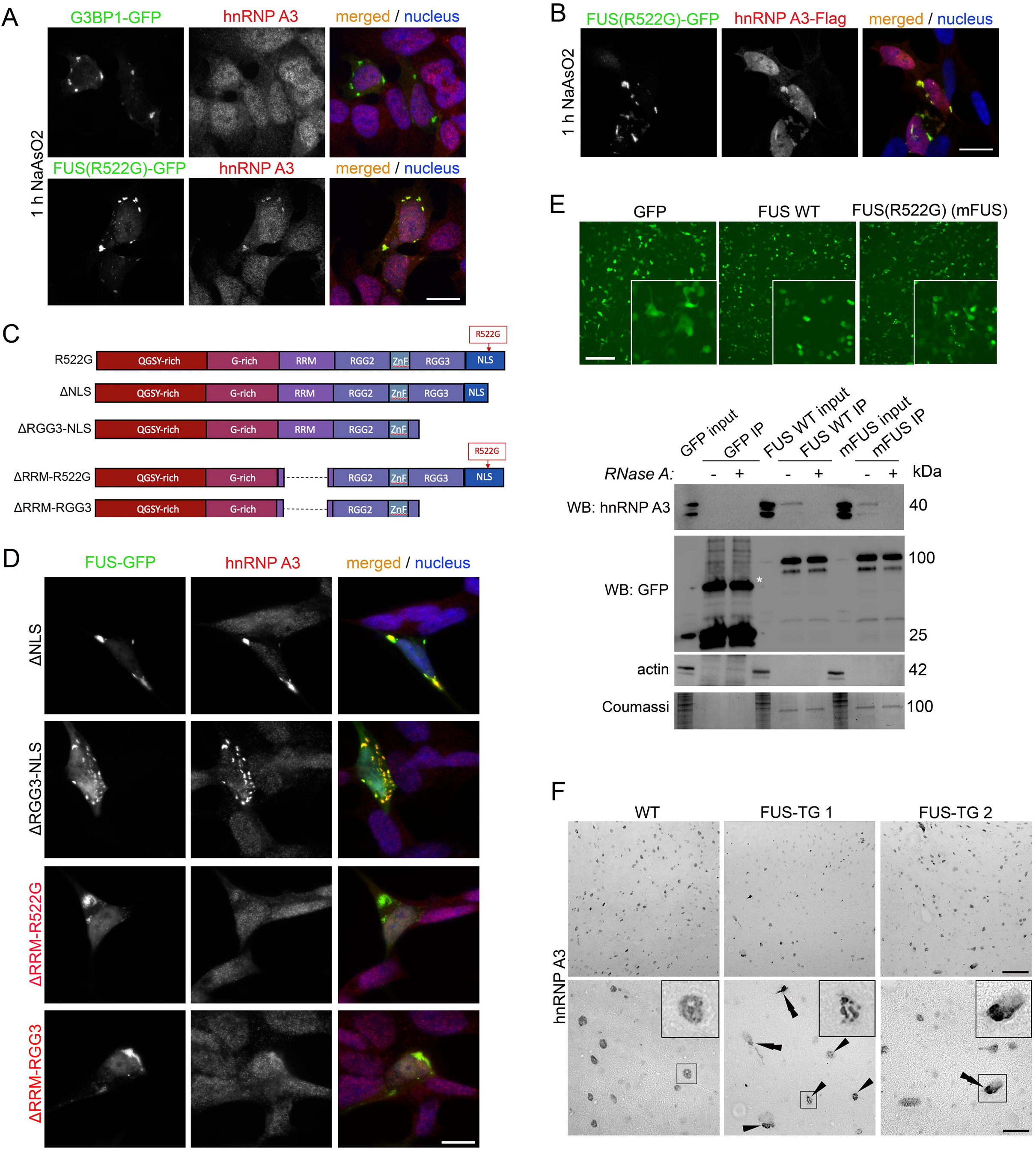
hnRNPA3 interacts with mutant FUS and is recruited into FUS aggregates in an RNA-dependent manner. (A) Enrichment of endogenous hnRNPA3 in mFAs formed by FUS(R522G)-GFP but not SGs formed by G3BP1-GFP in NaAsO_2_-treated cells. SH-SY5Y cells were treated with NaAsO_2_ for 1 h and immunostained with an anti-hnRNPA3 antibody. Scale bar, 10 µm. (B) Co-localization of overexpressed FUS(R522G)-GFP and hnRNPA3-Flag in mFAs in stressed SH-SY5Y cells. SH-SY5Y cells were treated with NaAsO_2_ for 1 h. Scale bar, 10 µm. (C) FUS deletion constructs used in the study. All FUS proteins were expressed as GFP fusions. (D) Mapping of FUS domains required for hnRNPA3 recruitment into mFAs. SH-SY5Y cells were analysed 24 h post-transfection with the respective construct, by anti-hnRNPA3 staining. FUS deletion mutants deficient in hnRNPA3 sequestration are given in red. Scale bar, 10 µm. (E) Co-IP of endogenous hnRNPA3 by GFP-tagged WT or mutant (R522G) FUS (mFUS) from overexpressing HEK293 cells. Top panel shows nuclear and cytoplasmic distribution of normal and mutant FUS, respectively, and efficiency of transfection in cultures used for co-IP (scale bar, 100 µm). Lysates of cells expressing GFP, FUS WT and mutant FUS were treated with RNase A or left untreated under the same conditions, and GFP-tagged proteins were precipitated using GFP-Trap® beads. Input is 10% of the final IP fraction. Asterisk indicates a non-specific band in “GFP only” pull-downs. (F) hnRNPA3 immunostaining of the spinal cord sections of WT and FUS-TG mice. Images for a 4-month old WT mouse and two symptomatic FUS-TG mice are shown. Nuclear inclusions are indicated with single arrowheads and cytoplasmic inclusions - with double arrowheads. Scale bar, 100 µm and 20 µm for general plane and magnified images, respectively.

We next examined the presence of hnRNPA3 in neuronal mutant FUS inclusions in a transgenic mouse model of FUS proteinopathy (Shelkovnikova et al., 2013a). These mice express a truncated version of FUS lacking RGG boxes, ZnF domain and therefore not expected to efficiently interact with hnRNPA3 (see Fig 4D). However inclusions formed by this variant also sequester normal endogenous FUS (Shelkovnikova et al., 2013a; Shelkovnikova et al., 2013b) which, as we hypothesized, may piggy-back hnRNPA3 into these structures. Immunostaining in the spinal cord tissue from symptomatic 4-month old mice showed accumulation of hnRNPA3 in a form of cytoplasmic and nuclear inclusions in neurons of transgenic mice, whereas WT mice displayed only normal nuclear hnRNPA3 staining (Fig. 4F).

Sequestration of hnRNPA3 into FUS aggregates/inclusions may lead to its loss of function. To examine whether and how downregulation of hnRNPA3 modifies FUS toxicity *in vivo*, we generated FUS WT transgenic *Drosophila* with silenced retinal expression of the fly hnRNPA3 ortholog, Hrb87F (Fig. 5A). As reported previously, FUS WT overexpression in the fly retina results in retinal thinning (∼30% reduced retinal thickness) (Matsukawa et al., 2021; Matsumoto et al., 2018) (Fig. 5A,B). We found that RNAi of Hrb87F also led to retinal thinning, comparable in its severity with that caused by FUS (Fig. 5A,B), pointing to an important housekeeping role of the hnRNPA3 ortholog in flies. However, in double transgenic flies, retinal degeneration was significantly more pronounced than in FUS WT or Hrb87F-RNAi flies, leading to almost complete loss of ommatidia (Fig. 5A,B). This result is indicative of additive toxicity of FUS accumulation and loss of Hrb87F expression. It should be noted that FUS levels were reduced in double transgenic flies as measured by western blot (Fig. 5C), which can be caused by severe retinal thinning in these flies or regulation of FUS levels by Hrb87F protein. Thus, loss of function of hnRNPA3 ortholog exacerbates FUS toxicity in flies.

**Figure 5.**
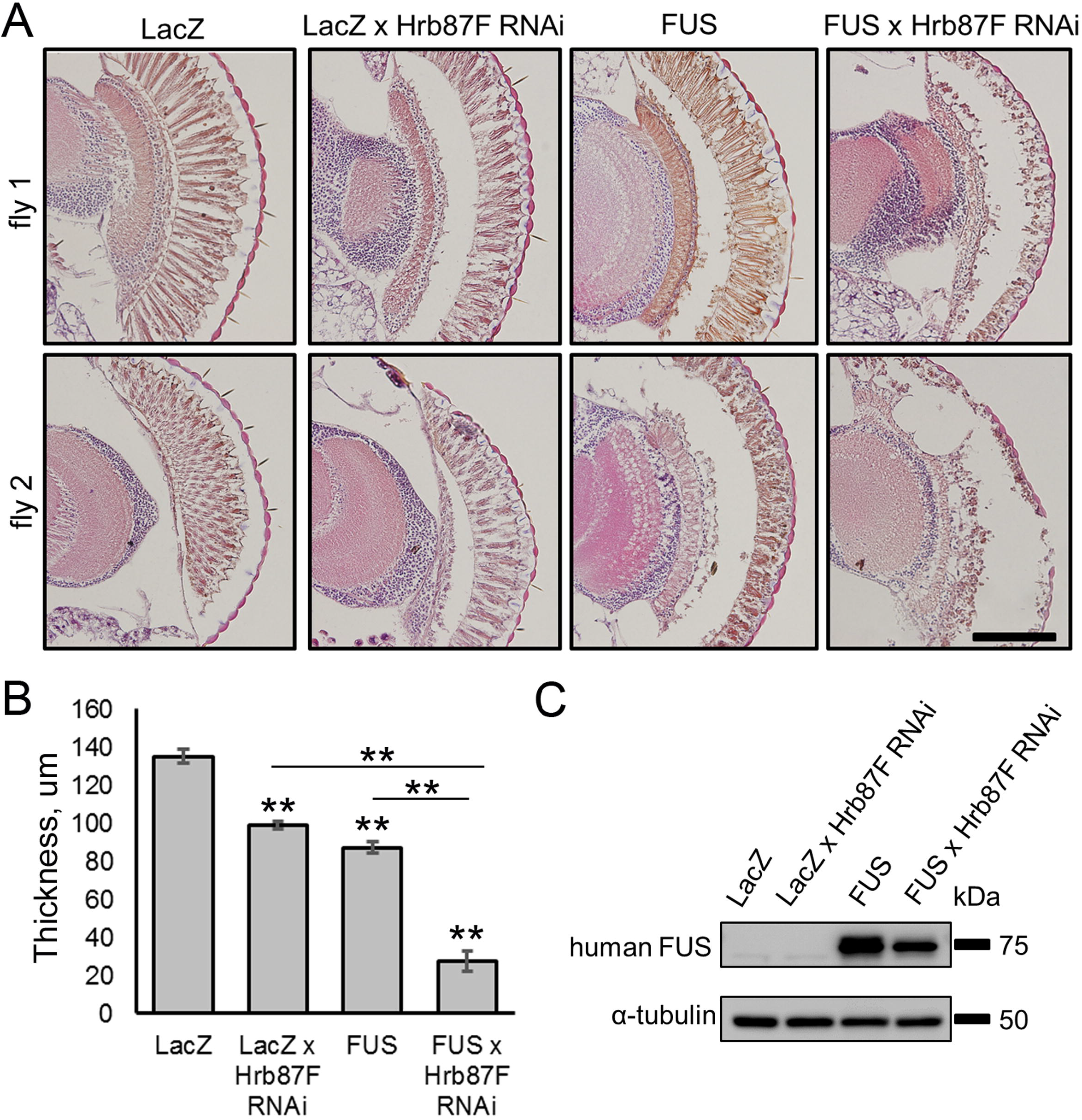
Depletion of the *Drosophila* hnRNPA3 ortholog exacerbates the toxicity of human FUS in the fly retina. (A,B) Downregulation of Hrb87F, the fly ortholog of human hnRNPA3, using RNAi, exacerbates FUS WT toxicity in the *Drosophila* retina. Representative images of retinal sections (A) and quantification of retinal thickness (B) are shown. In B, n=10 for each genotype, error bars represent S.E.M. **p<0.01 (one-way ANOVA with post-hoc Tukey-Krammer test). Scale bar, 100 µm. (C) Levels of FUS WT in the *Drosophila* retina in single and double (FUS/Hrb87F) transgenic flies. A representative western blot is shown.

## Discussion

In the current report, we provide proteomic and transcriptomic evidence, backed by cellular and *in vivo* validation studies, that: *i)* despite sharing many protein and RNA components, cytoplasmic RNP granules formed by mutant FUS in stressed cells are compositionally distinct from physiological SGs; *ii)* presence of cytoplasmic FUS assemblies is expected to significantly impact on the cellular protein and RNA homeostasis under stress. Our data strongly support a pathological role for stress-induced FUS aggregation in the cytoplasm in ALS-FUS, realized via its negative impact on SG function and gain of novel unwanted functions.

In a recent report, FUS aggregates were captured using a membrane filtration protocol with subsequent proteomic analysis (Kamelgarn et al., 2018). In another recent study, condensed FUS droplets from cell lysates were crosslinked, FACS-sorted and immunoprecipitated, with proteomic and transcriptomic analyses carried out on pulled WT/mutant FUS preparations separately for soluble and phase-separated fractions (Reber et al., 2019). These studies identified core cellular pathways regulated by FUS protein and confirmed that some of them, such as translation and NMD, are compromised in mutant FUS expressing cells. The former report found that mutant FUS is more deleterious for cellular processes than WT protein, whereas the latter provided evidence that mutant FUS toxicity might be partially dependent of its ability to phase-separate and aggregate. FUS phase separation and aggregation, including partitioning into SGs, is dramatically enhanced by stress. In addition, stress is believed to play an important role as a secondary trigger, or second hit, in ALS-FUS development (Al-Chalabi et al., 2014; Dormann et al., 2010; Shelkovnikova et al., 2019). Therefore it still remained to be addressed how mutant FUS assemblies relate to physiological stress-induced protein aggregation in the cytoplasm. FUS assemblies are often referred to as pathological SGs or SG-like structures, but until now, differences and similarities between these two types of RNP granules remained elusive. Indeed, FUS was shown to recruit some factors not normally present in physiological SGs, such as P-body, paraspeckle and spliceosome components (Gerbino et al., 2013; Shelkovnikova et al., 2014a; Shelkovnikova et al., 2014b), but the wider spectrum of protein/RNA species connected to these granules remained unknown. We adopted an approach different from the published studies – a parallel affinity purification of the two types of stress-induced granules, physiological SGs and mutant FUS assemblies, which enabled us to compare their effect on cellular proteostasis and RNA metabolism.

Proteomic analysis of SGs and mFAs showed that a certain “core” network of proteins related to the RNP granule assembly, regulation of protein translation, RNA metabolism and gene expression are still recruited into pathological FUS-containing RNP granules during stress. Therefore, mFAs may adopt some functions typical for physiological SGs. Yet, full functional replacement of SGs by mFAs is hardly possible because SG and mFA cores share only one-third of the proteome.

We found that two classes of proteins, NPC proteins and proteasome subunits are depleted from mFAs, as compared to SGs. Nucleocytoplasmic transport factor sequestration into SGs is likely an important component of cellular stress response, which when dysregulated, can contribute to ALS pathology (Zhang et al., 2018). Recently, 26S proteasome was shown to be recruited into arsenite-induced SGs to enable their clearance post-stress, whereas impaired proteasome function can lead to SG transformation into aberrant structures that require autophagic clearance (Turakhiya et al., 2018). Thus, insufficient NPC factor recruitment can impair stress signaling pathways, whereas lack of recruitment of the proteasome machinery into mFAs during stress might make them prone to persistence and subsequent conversion into insoluble aggregates. Recently, SGs have been shown to recruit sumoylation factors which facilitate their disassembly, and these factors are depleted from SGs formed in cells accumulating C9ORF72 pathological dipeptides (Marmor-Kollet et al., 2020). Interestingly, the SG but not mFA dataset in our study demonstrated enrichment for sumoylation proteins, which might contribute to impaired disassembly of FUS aggregates.

We found that several classes of proteins are abnormally enriched in mFAs, as compared to SGs, including mitochondrial proteins, translation and mRNA surveillance pathway factors, and components of the spliceosome, which is in line with reports by others. Enrichment of mFAs in mitochondrial proteins is consistent with a previous finding that FUS interacts with mitochondria (Deng et al., 2015), and a recent report of specific interaction of phase-separated FUS with mitochondrial proteins and entrapment of mitochondrial ribosomal RNAs in these aggregated FUS fractions (Reber et al., 2019). Binding of translation factors by mutant aggregated FUS was also recently demonstrated (Kamelgarn et al., 2018), and the effect of FUS aggregates on translation was highlighted in several studies (Kamelgarn et al., 2018; Lopez-Erauskin et al., 2018; Scekic-Zahirovic et al., 2016; Yasuda et al., 2013). Components of the CPSF complex (CPSF7, CPSF6 and CPSF1) and the nuclear cap-binding complex (CBC - NCBP1, NCBP2), were also identified in mFA cores but not in SG cores. mRNA surveillance pathway factors UPF1 and UPF3 were previously found to be dysregulated in cells containing mutant FUS aggregates and sequestered by aggregated FUS (Kamelgarn et al., 2018). In our dataset, UPF1 was present both in SG and mFA cores, although we could not estimate the efficiency of its recruitment quantitatively. UPF1 also belongs to the NCBP1 interaction network according to the STRING database. The CBC complex is essential for a pioneer round of mRNA translation, which also plays a central role in NMD. Mislocalization/dysregulation of spliceosome subunits in mutant FUS expressing cells (Gerbino et al., 2013; Reber et al., 2016) and the negative effect of mutant FUS on nuclear SMN-containing RNP granules, Gems (Yamazaki et al., 2012) have been established. Abnormal sequestration of proteins involved in major (U2) and minor (U12) spliceosome subunits as well as Gemins into mFAs is in line with this reported role of FUS in the spliceosome. Interestingly, spliceosome and mRNA surveillance factors DAZAP1 and CPSF6 sequestered into mFAs are also components of nuclear RNP granules paraspeckles. DAZAP1 is a core paraspeckle protein responsible for the stability of this granule, and loss of function of CPSF6 is known to lead to enhanced accumulation of the structural paraspeckle RNA, NEAT1_2 (Naganuma et al., 2012). Recently we demonstrated that paraspeckle integrity is affected in cells expressing mutant FUS despite accumulation of NEAT1_2 (An et al., 2019a).

At the RNA level, we found that overall, mFA-recruited transcripts displayed a less significant enrichment as compared to SGs. This result may suggest less efficient separation of mFAs from the surrounding cytosol. We found that although mFAs recruit the RNA species generally mapping to the same processes and pathways (e.g. transcription and protein phosphorylation) as those recruited to SGs, the diversity of these transcripts is significantly decreased. Regulation of specific functional sets of transcripts can be attenuated in the presence of mFAs that disrupt SGs, where SG function in sequestration of these RNAs is partially lost. Loss of sequestration for these transcripts may lead to their enhanced degradation or their unlicensed translation during stress. In particular, we found that recruitment of mRNAs encoding stress-activated MAPK cascade, mTOR signaling, regulation of protein translation and apoptotic processes was affected, which may lead to suboptimal responses during recovery from stress and impact cell survival.

Our data suggest that the majority of pathways dysregulated by mutant FUS under basal conditions, e.g. protein translation, RNA quality control, mitochondrial function, splicing, DNA damage repair and chromatin remodeling (Kamelgarn et al., 2018; Reber et al., 2019), are also affected under stress. However, our approach allowed identification of new pathways and factors dysregulated by mutant FUS assemblies at the protein and RNA level specifically under stress, first of all, proteasome clearance of misfolded proteins/aggregates (at the protein level) and cellular stress/apoptotic signaling (at the RNA level). Notably, while SGs form in cells only in response to acute stress, FUS aggregates can form spontaneously and persist for prolonged periods of time. They therefore can induce a stress-mimicking state in neurons by retention of proteins and RNAs normally sequestered into SGs. This “preconditioned”, chronic stress state might render neurons more vulnerable under acute stress conditions (Shelkovnikova et al., 2017).

A hit from our proteomic analysis, hnRNPA3, was identified as a modifier of FUS toxicity *in vivo*, in transgenic flies. Loss of hnRNPA3 function caused by its entrapment in FUS inclusions is expected to worsen the toxic effect of FUS, similar to the negative effect of hnRNPA3 loss of function in ALS-C9 (Mori et al., 2013; Nihei et al., 2020). In the latter ALS subtype, this effect was shown to be mediated by altered DNA damage response, whereas specific effects in ALS-FUS are yet to be established. hnRNAP3 has also been found significantly downregulated in cultured cells after knockdown of another major ALS-associated protein, TDP-43, whose aggregation is typical for ALS-C9 (Prpar Mihevc et al., 2016). Interestingly, the fly ortholog of hnRNPA3, Hrb87F, has been identified as an enhancer of TDP-43 toxicity (Appocher et al., 2017). Therefore, the role of hnRNPA3 might be different in the context of ALS-FUS and ALS-TDP pathogenesis.

In conclusion, this research lays the groundwork for studies of other ALS-causative mutants – SG components in cellular stress response. Cataloguing differences between physiological SGs and their abnormal, disease-associated counterparts should allow the establishment of common pathways and factors in genetically different ALS cases and therefore inform on universal therapeutic targets in ALS.

## Figure legends

**Figure S1. Comparisons of HEK293 proteomic and transcriptomic SG**_**core**_ **datasets with the published datasets for U2OS cells**.

(A) Overlaps in Biological Process GO term between HEK293 (this study) and U2OS (Jain et al., 2016) SG_core_ proteomic datasets.

(B) Biological Process GO terms enriched in HEK293 (this study) and U2OS SG_core_ (Jain et al., 2016) proteomic datasets.

(C) Overlaps in Biological Process GO terms between the HEK293 (this study) and U2OS (Khong et al., 2017) SG_core_ transcriptomic datasets.

(D) Biological Process GO terms enriched in the HEK293 (this study) and U2OS (Khong et al., 2017) SG_core_ transcriptomic datasets.

## Materials and methods

### Cell culture and maintenance

HEK293 and SH-SY5Y cells were maintained in DMEM/F12 medium supplemented with 10% foetal bovine serum (FBS), penicillin-streptomycin and GlutaMAX® (all Invitrogen). Production of plasmids encoding G3BP1-GFP (pEGFP-N1 vector), FUS(R522G)-GFP (pEGFP-C1 vector) and FUS deletion mutants is described in our previous studies (An et al., 2019b; Shelkovnikova et al., 2014a). Plasmid for the expression of Flag-tagged hnRNPA3 was purchased from Sino Biological (Cat #CG90366-NF-SIB). For small-scale transfections in cellular validation analysis, Lipofectamine2000 (Invitrogen) was used in 24-well plates. Cells were treated with 0.5 mM NaAsO_2_ (sodium arsenite) for 1 h to induce SG and mFA assembly.

### Stress granule core and FUS granule affinity purification

SG and mFA cores were purified from HEK293 cells transiently expressing G3BP1-GFP or FUS(R522G)-GFP according to a previously published protocol, with modifications (Jain et al., 2016; Wheeler et al., 2017). HEK293 cells were transfected with plasmids to express GFP alone, G3BP1-GFP and FUS(R522G)-GFP in 6-cm dishes (1 µg plasmid/dish) using Lipofectamine2000. The following day (∼24 h post-transfection), cells were treated with 0.5 mM NaAsO_2_ for 1 h, snap-frozen on dishes and processed for SG/mFA core purification in parallel. Cells were scraped in the lysis buffer containing 50 mM TrisHCl pH 7.4, 100 mM KOAc, 2 mM MgOAc, 0.5 mM DTT, 50 µg/ml heparin, 0.5% NP-40 and supplemented with RNase inhibitor (Murine, M0314, New England Biolabls) and protease inhibitors cocktail (cOmplete Mini, Roche), and passed through a 25G needle 7 times. Lysates were left on ice for 15 min with periodic vortexing. 1/50 of the lysates was kept for preparation of total cellular RNA for transcriptomic analysis. Lysates were subsequently centrifuged at 1,000xg for 5 min, and the supernatant was centrifuged again at 17,000xg for 20 min, to pellet the cores. The pellets washed twice in the lysis buffer, resuspended in the same buffer and incubated with GFP-Trap® agarose beads for 4 h. Beads were washed three times with washing buffer 1 (20 mM TrisHCl, 200 mM NaCl, pH 8.0), once with washing buffer 2 (20 mM TrisHCl, 500 mM NaCl, pH 8.0) and once with washing buffer 3 (lysis buffer supplemented with 2M urea). 1/8 of the resultant bead slurry was used directly for proteomic (LC-MS/MS) analysis and the remaining beads were used for RNA purification. Purification of SG and mFA cores was performed in duplicates and twice, on two different days, and proteomic and transcriptomic analyses was done on two samples (combination of two biological replicates each) per condition.

### Proteomic analysis

Proteomic analysis was performed at the Bristol Proteomics Facility as described in (An et al., 2019b). Peptide data were filtered to satisfy false discovery rate (FDR) of 1%. Peptides identified were mapped to proteins using the respective tool of the UniProt online database (https://www.uniprot.org/). Proteins identified in the samples from cells expressing GFP alone were used as a background list and were subsequently subtracted from final SG and mFA core protein lists. Venn diagrams were prepared using BioVenn online tool (http://www.biovenn.nl/index.php) (Hulsen et al., 2008). Protein networks were prepared using the STRING Database v.11 (https://string-db.org/). Enrichment analysis was carried out using Enrichr free online tool (https://amp.pharm.mssm.edu/Enrichr/) (Chen et al., 2013; Kuleshov et al., 2016) and Revigo (http://revigo.irb.hr/) (Supek et al., 2011).

### RNA analysis

RNA was purified from lysates (total cellular RNA) and separately from SG or mFA core fractions using TRI-reagent (Sigma). Total RNA was approximately 10 times more concentrated than RNA from SG/mFA core fractions (∼100 ng/µl *vs*. 10 ng/µl) and therefore was diluted accordingly prior to RNA-Seq analysis. For validation/quality control experiments, cDNA synthesis was performed using random primers (Promega) and M-MLV reverse transcriptase (Promega), according to manufacturer’s instructions. Non-saturated PCR (26 cycles) was performed using New England BioLabs Taq DNA polymerase (M0273). Primer sequences were: CHD7: 5’-GCAGAAAGTGCCTGTGCATC-3’ and 5’-GCTGAGCATTCGGTCCACTA-3’; BCL9L: 5’-CGTACAGTGGGGACGAATGG-3’ and 5’-ATGGCTGGGTCTGCTACATT-3’; UBL5: 5’-CTGATTGCAGCCCAAACTGG-3’ and 5’-CAGGTTCATCCCATCGTGGA-3’; MACF: 5’-TGCATGAGCAGAAAAAGCGG-3’; 5’-TTTCTTCTGAACCCGGTCCC-3’; RPL39: 5’-GTGTGTTCTTGACTCCGCTG-3’ and 5’-TTCATCCGAATCCACTGGGG-3’; GAPDH: 5’-TCGCCAGCCGAGCCA-3’ and GAGTTAAAAGCAGCCCTGGTG-3’. RNA-Seq analysis was performed at the School of Biosciences Genomics Research Hub. Libraries were prepared using the TruSeq stranded total kit (Illumina), and single-end sequencing was performed on Illumina NextSeq500 (read length: 75_D_bp; coverage ∼_D_50 million reads/sample). Reads were aligned to GRCh38 human genome using STAR aligner and Gencode release 31 annotation. Gene level quantification was also obtained using STAR (Dobin et al., 2013). To calculate enrichment and depletion of RNAs in SG and mFA cores, differential gene expression analysis using the voom-limma method was applied (Law et al., 2014). This analysis was performed for pairs “SG total lysate – SG core” and “mFA total lysate– mFA core”. Genes “upregulated” and “downregulated” in SG or mFA cores were considered enriched and depleted, respectively. Out of 18,068 transcripts identified, 1,645 were enriched in SG cores and 3,279 were depleted from SG cores; and out of 18,416 transcripts identified, 802 were enriched in mFA cores and 2,431 were depleted from mFA cores. Fold-change (FC) cut-off was 1.2 and p-value cut-off was 0.5 both for enriched and depleted RNAs. Enrichment analysis using Enrichr / Revigo was performed as described for proteomics.

### Immunocytochemistry

Immunocytochemistry and microscopic analysis were performed as described earlier (Kukharsky et al., 2015), with modifications. Briefly, HEK293 or SH-SY5Y cells were plated on uncoated coverslips, transfected and stressed 24 h post-transfection, fixed 4% paraformaldehyde for 15 min at RT and permeabilized for 5 min in cold methanol. Primary antibodies in blocking solution (5% goat serum in 0.1% Triton-×100/1xPBS) were applied for 2 h at RT or, when needed, overnight at 4°C. Secondary Alexa488- or Alexa546-conjugated antibodies (Molecular Probes) separately or in cocktail were added for 1 h at RT. Nuclei were visualised with a 5 min incubation in 10 µg/ml DAPI solution (Sigma). Fluorescent images were taken with 100x objective (UPlanFI 100x/1.30) on BX57 fluorescent microscope equipped with ORCA-Flash 4.0 camera (Hamamatsu) and cellSens Dimension software (Olympus). Figures were prepared using Photoshop CS3 or PowerPoint 2016 software.

### Immunoprecipitation (IP) and western blotting

Cells were washed with PBS, lysed in ice cold IP buffer (1% Triton-×100 in PBS) on ice with periodic vortexing for 15 min. Unbroken cells and cell debris were pelleted at 13,000 rpm for 15 min, and input samples were taken at this point. Cleared cell lysates were split in half and one half was treated with RNase A (100 µg/ml) for 30 min at RT. Lysates were then incubated with GFP-Trap® agarose beads (ChromoTek) for 4 h at 4□C. Beads were washed four times with ice cold IP buffer, and bound protein complexes were eluted by heating the samples for 10 min at 95□C in 2xLaemmli loading buffer. Pull-down efficiency was analysed by western blot. For input, 10% of the final IP sample was loaded on the gel. Proteins were resolved in Mini-Protean® TGX precast gels (Bio-Rad) and transferred to the PVDF membrane (GE Healthcare) by semi-dry blotting. Membranes were blocked in non-fat 4% milk in TBS/T and incubated with primary and HRP-conjugated secondary (GE Healthcare) antibodies. For signal detection, Clarity Max ECL kit and ChemiDoc™ Gel Imaging System (Bio-Rad) were used.

### Primary antibodies

The following commercial primary antibodies were used: hnRNPA3 (rabbit polyclonal, 25142-1-AP, Proteintech); CPSF6 (rabbit polyclonal, A301-356A, Bethyl); NCBP1 (rabbit polyclonal, 10349-1-AP, Proteintech); DAZAP1 (rabbit polyclonal, A303-984A, Bethyl); GFP (mouse monoclonal, sc-9996, Santa Cruz); EXOSC3 (rabbit polyclonal, 15062-1-AP, Proteintech); Flag (DYKDDDDK Tag, mouse monoclonal, 9A3, Cell Signaling); beta-actin (mouse monoclonal, A5441, Sigma). Antibodies were used at 1:1,000 dilution for immunostaining and western blot.

### Immunohistochemistry on mouse samples

Spinal cord sections (8 µm thick) from wild-type and 4-month old symptomatic FUS-TG mice (Shelkovnikova et al., 2013a) were used. After rehydration, sections were subjected to microwave antigen retrieval in sodium citrate buffer (pH 6.0) and blocked using 10% goat serum in PBS/T. Sections were incubated with the primary anti-hnRNPA3 antibody (Proteintech, 25142-1-AP) overnight at 4□C and secondary HRP-conjugated anti-rabbit IgG antibody (Vector Laboratories) for 1.5 h at RT. Signal was detected using Vectastain® Elite ABC Universal Plus Kit (Vector Laboratories) and 3,3′-diaminobenzidine (DAB, Sigma). Images were taken using BX57 microscope (Olympus) and ORCA-Flash 4.0 camera (Hamamatsu).

### Generation and analysis of transgenic Drosophila

Generation of *Drosophila* lines with human retinal expression of FUS WT is described elsewhere (Matsumoto et al., 2018). gmr-GAL4, UAS-lacZ and Hrb87F RNAi lines were obtained from the Bloomington Drosophila stock center. For external surface analysis of the fly eye, 5-day-old flies were anesthetised with CO_2_ and examined by zoom stereo microscopy (Olympus SZ-PT). For histochemical analysis, heads of 5-day-old adult flies were dissected, briefly washed in PBS and fixed with 4% PFA containing 0.1% Triton X-100 at RT for 2 h. After brief wash in PBS, tissues were dehydrated by graded ethanol, cleared in butanol and embedded in paraffin. Four-micrometre thick coronal sections were stained with hematoxylin and eosin (H&E). Ten animals of each genotype were used for retinal thickness quantification. For western blot analysis, heads of 5-day-old flies were dissected and lysed in Laemmli sample buffer for SDS-PAGE. Anti-FUS antibody from Bethyl (A300-293A) was used for western blot.

## Supporting information

Table S1. Proteomic datasets and their analysis

Table S2. Transcriptomic datasets and their analysis.

