## Supplementary figures and images for "Compositional analysis of ALS-linked stress granule-like structures reveals factors and cellular pathways dysregulated by mutant FUS under stress"

### Figure S1

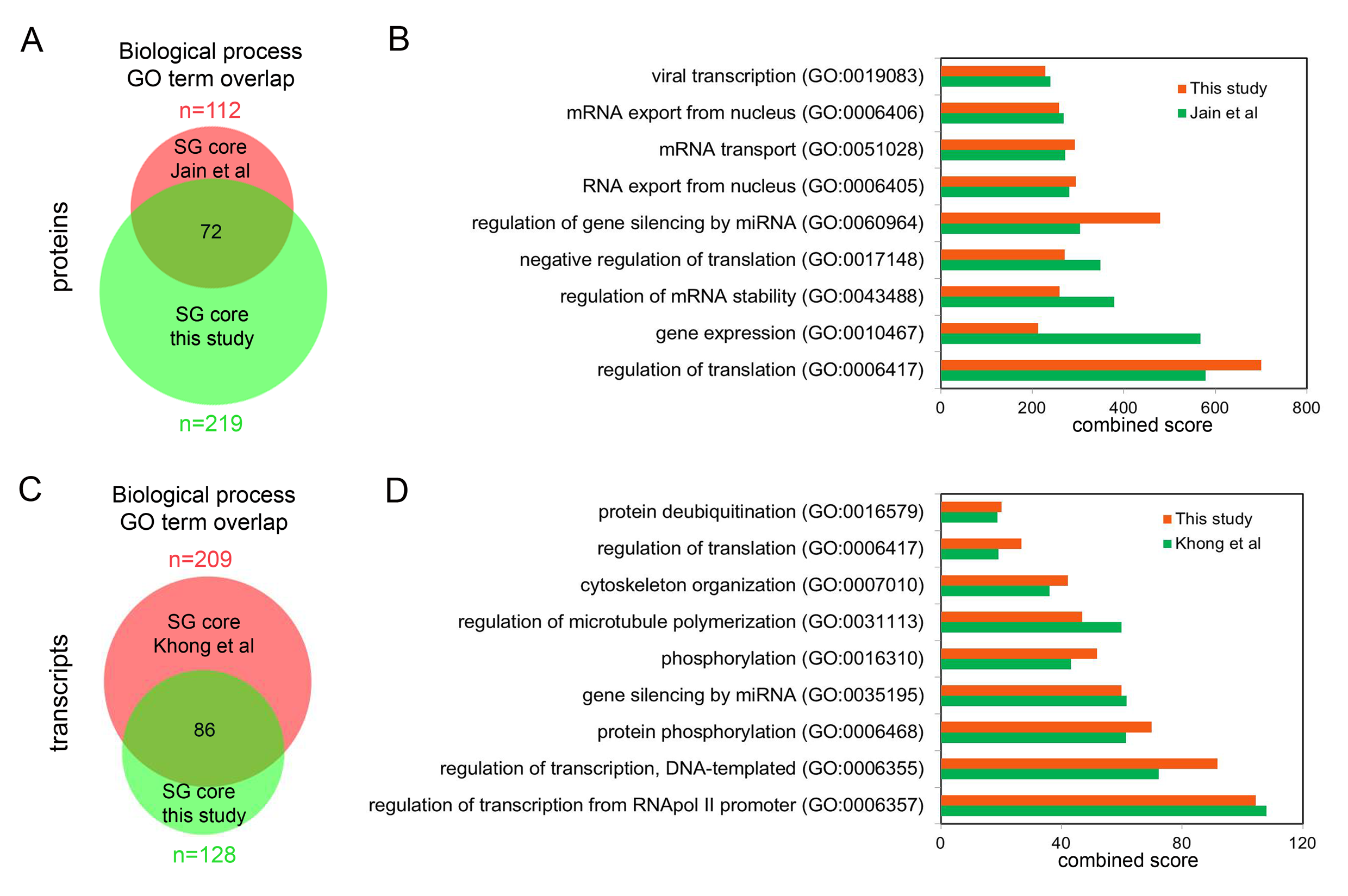
